# Microbiome signatures in a fast and slow progressing gastric cancer murine model and their contribution to gastric carcinogenesis

**DOI:** 10.1101/2020.10.15.341701

**Authors:** Prerna Bali, Joanna Coker, Ivonne Lozano-Pope, Karsten Zengler, Marygorret Obonyo

## Abstract

Gastric cancer is the third most common cancer in the world and *Helicobacter spp*. being one of the main factors responsible for development of cancer. Alongside *Helicobacter* the microbiota of the stomach mucosa may also play an important role in gastric cancer progression. Previously we had established that MyD88 deficient mice rapidly progressed to neoplasia when infected with *H. felis*. Thus, in order to assess the role of microbiota in gastric cancer progression we measured the changes in microbial diversity of the stomach in mice with different genotypic backgrounds (Wild type (WT), MyD88 deficient (MyD88^−/−^), mice deficient in the Toll/IL-1R (TIR) domain-containing adaptor-inducing interferon-β (TRIF, *Trif*^lps2^), and MyD88 and Trif deficient (MyD88^−/−^ and Trif^−/−^)double knockout (DKO) mice), both in uninfected and *Helicobacter* infected mice and its correlation of these changes with gastric cancer progression. We observed that there was an overall reduction in microbial diversity post infection with *H. felis* across all genotypes. *Campylobacterales* were observed in all infected mice, with marked reduction in abundance at 3 and 6 months in MyD88^−/−^ mice. This low abundance of *H. pylori* could facilitate dominance of other organisms of microbiome like *Lactobacilliales*. A sharp increase in *Lactobacilliales* in infected MyD88^−/−^ and DKO mice at 3 and 6 months was observed as compared to Trif^−/−^ and WT mice suggesting its possible role in gastric cancer progression. This was further reinforced upon comparison of *Lactobacillus* ratio with histological data suggesting that *Lactobacillales* is closely associated with *Helicobacter* infection and gastric cancer progression. Thus, this study firstly suggests that difference in genotypes could define the stomach microbiome and make it more susceptible to development of gastric cancer upon *Helicobacter* infections. Secondly the increase in *Lactobacillales* could contribute to faster development of gastric cancer and serve as a probable bio marker for fast progressing form of gastric cancer.

## Introduction

Gastric cancer is the sixth most common cancer and third most common in the world (1). By far the largest risk factor for gastric cancer development is the carcinogenic microbe *Helicobacter pylori* (2). Infection with *H. pylori* leads to development of premalignant lesions – that progress from gastric atrophy to metaplasia, dysplasia, and finally to gastric adenocarcinoma. *H. pylori* infects almost 50% of the global population but only 1-3% of the infected individuals develop gastric cancer (3).

Several other factors contribute to gastric cancer progression such as *Helicobacter* strains, environmental factors, and host immune response. In addition, the microbiota of the stomach may also influence the final disease outcome (3).

The stomach had been traditionally considered a sterile organ due to its highly acidic environment and digestive juices. It was only after the discovery of *H. pylori* in 1982, and with its ability to survive in harsh conditions that led to the idea that other microbiomes could inhabit the stomach. Advancement in DNA sequencing strategies and computational methods a complex microbiome of the stomach was has been uncovered (3).

The gastric microbial density is estimated to be 10^2^ - 10^4^ colony forming units (CFU)/ml, a comparatively lower density than the colon, which often reaches 10^10^ to 10^12^ CFU/ml (4). Early characterization of gastric microbiota relied on culturing techniques but with the advancement of sequencing techniques led to the identification of different species falling into five predominant phyla Firmicutes, Proteobacteria, Actinobacteria, Bacteroidetes and Fusobacteria, indicating a distinct microbiome of the stomach (4,5). Thus, with the presence of such complex microbial diversity in the stomach interactions between *Helicobacter* and other gut bacteria could play a role in deciding the fate of progression of gastric cancer or may act as an aide along with other factors in driving the disease.

We have previously shown that Myeloid differentiation primary response gene 88 (MyD88) regulates *Helicobacter* induced gastric cancer progression, where Myd88 knockout (MyD88^−/−^) mice infected with *Helicobacter felis* showed fast progression to adenocarcinoma as compared to wild type (WT) mice (6).

This prompted us to study the microbial diversity of the stomach in mice with different genotypic backgrounds (Wild type (WT), MyD88 deficient (MyD88^−/−^), mice deficient in the Toll/IL-1R (TIR) domain-containing adaptor-inducing interferon-β (TRIF, *Trif*^lps2^), and MyD88 and Trif deficient (MyD88^−/−^ and Trif^−/−^)double knockout (DKO) mice), both in uninfected and *Helicobacter* infected mice. In addition, we also investigated whether the gastric microbiome changes over time in response to infection and studied the correlation of these changes with gastric cancer progression. We hypothesize that this would help us in identifying, the microbial species whose abundance or scarcity may contribute to progression of *Helicobacter* induced lesions, either rapidly or slowly towards adenocarcinoma, that could open new avenues of research for gastric cancer.

## Material and Methods

### Animals

Six- to ten- week-old wild type (WT) (n=42), MyD88 deficient (Myd88^−/−^) (n=47), Trif deficient (Trif^−/−^) (n=46), Double knockout (MyD88^−/−^ Trif ^−/−^) (n=37) mice in the C57BL/6 background were used in this study. WT mice were purchased from The Jackson Laboratory (Bar Harbor, ME). Myd88^−/−^ mice were from our breeding colony originally provided by Dr. Akira (Osaka University, Japan). Trif^−/-^ mice and Double knockout mice were from our in-house breeding colony. All mice were housed together before infection with *H. felis* and for the duration of the study for each genotype. The institutional Animal Care and Use Committee at the University of California, San Diego, approved all animal procedures and performed using accepted veterinary standards.

### Bacterial Growth Conditions

*Helicobacter felis*, strain CS1 (ATCC 49179) was purchased from American Type Culture collection (Manassas, VA). *H. felis* was routinely maintained on solid medium, Columbia agar (Becton Dickinson, MD) supplemented with 5% laked blood under microaerophilic conditions (5% O2, 10% CO2, 85% N2) at 37 °C and passaged every 2–3 days as described previously (6-8). Prior to mouse infections, *H. felis* was cultured in liquid medium, brain heart infusion broth (BHI, Becton Dickinson) supplemented with 10% fetal calf serum and incubated at 37 °C under microaerophilic conditions for 48 h. Spiral bacteria were enumerated using a Petroff-Hausser chamber before infections.

### Mouse Infections

A well-characterized cancer mouse model, which involves infecting C57BL/6 mice with *H. felis* (strain CS1), a close relative of the human gastric pathogen *H. pylori* was used in this study. Mice were inoculated with 10^9^- organisms in 300 μL of BHI by oral gavage three times at 2-day intervals as previously described (6,7). Control mice received BHI only. At 1 month, 3 months and 6 months post infection, mice were euthanized, and the stomachs removed under aseptic conditions. The stomach was cut longitudinally and tissue sections were processed for DNA extraction and histopathology.

### Histology

Longitudinal sections of stomach tissue from each mouse were fixed in neutral buffered 10% formalin and embedded in paraffin, and 5-μm sections were stained with hematoxylin and eosin (H&E). Gastric histopathology mucous metaplasia, was scored by a blinded comparative pathologist (Rickman) using criteria developed by Rogers et al. (9). Scores ranging from 0 (no lesions) to 4 (severe lesions) were measured in increments of 0.5, as previously described.

### DNA Extraction

DNA was extracted from gastric tissue obtained from *H. felis*-infected and uninfected WT, Myd88^−/−^ KO, Trif^−/−^ KO, and MyD88^−/−^ Trif ^−/−^ Double knockout mice. Stomach tissue sections were analyzed at different time points of 1 month, 3 months, and 6 months. DNA was extracted using DNAeasy Blood &Tissue kit (Qiagen) following manufacturer’s instructions. The DNA concentration was quantified using NanoDrop 1000 Spectrophotometer (Thermo Fisher Scientific, Waltham, MA, USA) before proceeding to DNA Sequencing.

### 16S rRNA gene sequencing

Purified DNA was amplified and processed according to Earth Microbiome Project (EMP) standard protocols (https://www.earthmicrobiome.org/protocols-and-standards/16s/) Amplicon PCR was performed on the V4 region of the 16S rRNA gene using the primer pair 515F/806R with Golay error-correcting barcodes on the reverse primer. 240 nanograms of each amplicon was pooled and purified with the MoBio UltraClean PCR cleanup kit (Qiagen) and sequenced on the Illumina MiSeq sequencing platform.

Demultiplexed fastq files were processed using QIIME2 (https://qiime2.org) (10). Sequences were denoised using Deblur (11). Taxonomy was assigned using SEPP fragment insertion with a classifier trained on the Greengenes13_8 99% OTUs dataset, with sequences trimmed to contain 250 bases from the region amplified in sequencing (12).

### Bioinformatics processing

Taxonomy abundance plots were generated using the PhyloSeq package (13). Taxonomy was collapsed to the order of interest in PhyloSeq. Alpha diversity plots were generated in R using data exported from the QIIME2 analysis. Beta diversity principal component analysis plots were generated in QIIME2 with the DEICODE plug-in (reference: C. Martino et al., A Novel Sparse Compositional Technique Reveals Microbial Perturbations. mSystems. 4 (2019), doi:10.1128/mSystems.00016-19). Taxonomy abundance log-fold differentials analysis was conducted using the Songbird plug-in (14), with visualization through Qurro (15). ROC curves were generated in Prism 7 using log-fold differential values from the Songbird analysis.

### Statistics

Differences in alpha diversity and log-fold differentials were calculated in R using Student’s t-test or ANOVA, where appropriate. Data was checked for normal distribution before testing. Correlation was assessed with Spearman’s rank-correlation. Differences in beta diversity were assessed with PERMANOVA in QIIME2 with Benjamini-Hochberg FDR correction.

## Results

### Gastric mucosal microbial diversity varies with infection status and in different genotypes

To assess the diversity of each gastric microbial community, we examined the alpha and beta diversity. The Shannon diversity index and Pielou’s evenness score were calculated as metrics of alpha diversity. The Shannon diversity index provides a metric of community diversity based on the number of taxa present and the abundance of each species. Differences in genotype did not significantly affect Shannon diversity index (Figure 1). In general, infection with *H. felis* resulted in a decreased diversity index (statistically significant for WT at 6 months (p<0.05); MyD88−/− at 1 month (p<0.05) and 3 months (p<0.01); Trif^−/−^ at 3 months (p<0.05); and double knockout at 3 months (p<0.01)). For the three knockout genotypes, diversity was decreased in infected samples at 1 and 3 months but increased back to uninfected levels at 6 months. The opposite was observed in the WT genotype. Pielou’s evenness assesses how evenly distributed taxa are within a community. The evenness scores follow the same trends as the Shannon diversity index, as expected (Figure 1).

**Figure 1.**
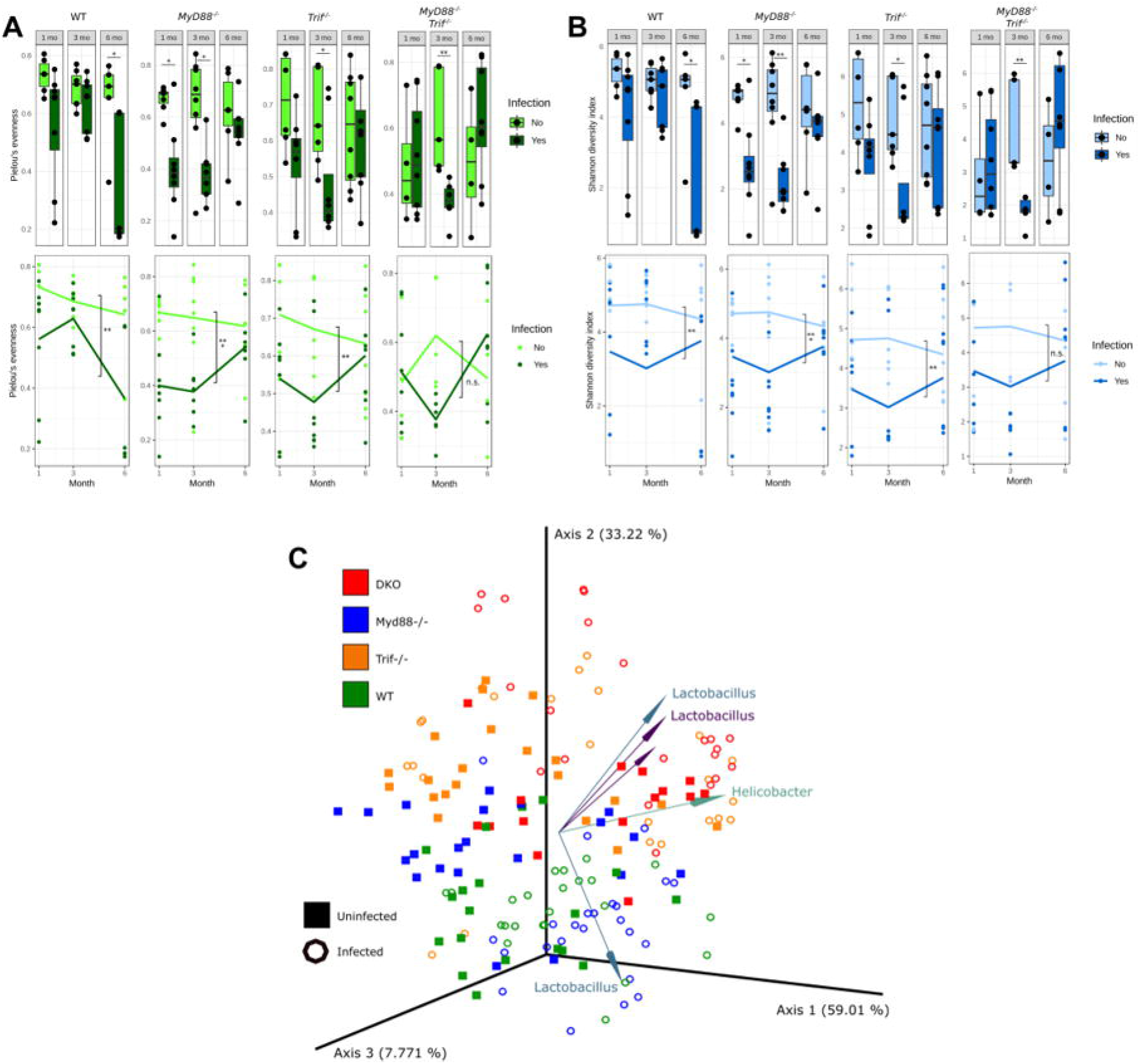
Changes in microbial diversity in the stomach across four different genotypes, with infection and time. Pielou’s evenness (A) and Shannon diversity index (B) values for each gastric community, broken out by genotype and month. Top and bottom plots represent the same data, with lines on the bottom plots representing average values and infected and uninfected communities over time. C) PCA of robust Aitchison distance values between communities. Biplot arrows indicate operational taxonomic units (OTUs) driving separation between categories, with arrows labeled with the genus of the OTU. Data is from months 1-6. All diversity metrics were calculated using QIIME2. Statistical significance determined by student’s t-test (* p<0.05, ** p<0.01, *** p<0.005).

To assess the dissimilarity between the gastric microbial communities, we next analyzed the robust Aitchison distance between communities. Aitchison distance is a compositional metric of the Euclidean distance between samples after centered log-ratio transformation (16). Robust Aitchison distance analysis incorporates matrix completion to account for the large number of zeros in microbiome data sets due to the absence of individual taxa in samples (17). PERMANOVA analysis of the robust Aitchison distance between samples showed significant separation between communities by all genotype pairs (p<0.001) except WT and Myd88^−/−^ (p=0.20). Uninfected and infected communities were significantly different within each genotype (p<0.001), despite continuing to cluster roughly with uninfected samples of the same genotype. Additionally, pairwise comparison (with Benjamini-Hochberg FDR correction) showed that infection groups within each genotype were significantly different from other groups (p<0.05), with the exception of WT uninfected/infected (p=0.07), MyD88^−/−^ uninfected/infected (p=0.07), WT infected/MyD88^−/−^ infected (p=0.15), Trif^−/−^ infected/MyD88^−/−^Trif^−/−^ uninfected (p=0.42), and Trif^−/−^ infected/MyD88^−/−^Trif^−/−^ infected (p=0.05). Principal coordinate analysis displayed that WT/MyD88^−/−^ genotypes and Trif^−/−^/DKO genotypes have more similar gastric communities, despite statistically significant differences (Figure 1). A biplot of the robust Aitchison PCA further revealed that the operational taxonomic units most heavily influencing the distance between samples are from the *Helicobacter* and *Lactobacillus* genera.

### Variation in abundance of microbial taxa in different genotypes over time and after *H. felis* infections

Given the differences in community diversity between genotypes and infection status, we conducted taxonomic analysis of the 16S rRNA gene sequences, 4 phyla, *Bacteroidetes*, *Cyanobacteria*, *Firmicutes* and *Proteobacteria* were predominant across all genotypes, irrespective of infection status and time (Figure 2). *Bacteroidetes* and *Firmicutes* were present both in uninfected and infected samples with insignificant variations between genotypes. *Cyanobacteria* was observed to be present in uninfected samples of all the four genotypes but more predominant in MyD88^−/−^ and DKO uninfected mice (Supplementary Figure 1). However, in stark contrast *Cyanobacteria* was completely absent in MyD88^−/−^ infected mice and its level dropped significantly in DKO infected samples at 1 month and were absent at both 3 and 6 months. On the other hand, *Cyanobacteria* levels were more predominant in infected Trif^−/−^ mice at 1 month and 3 months. *Proteobacteria* was observed in high abundance in infected samples as compared to uninfected samples in all genotypes. However, the levels of *Proteobacteria* dropped significantly at 3 months and 6 months in infected MyD88^−/−^ mice and at 6 months for DKO mice in infected samples. In contrast, in Trif^−/−^ knockout and wild type (WT) no such drop is observed. Moreover, the levels of *Proteobacteria* in Trif^−/−^ knockout were similar at 1 month and 6 months, while peaking at 3 months (Supplementary Figure 1).

**Figure 2.**
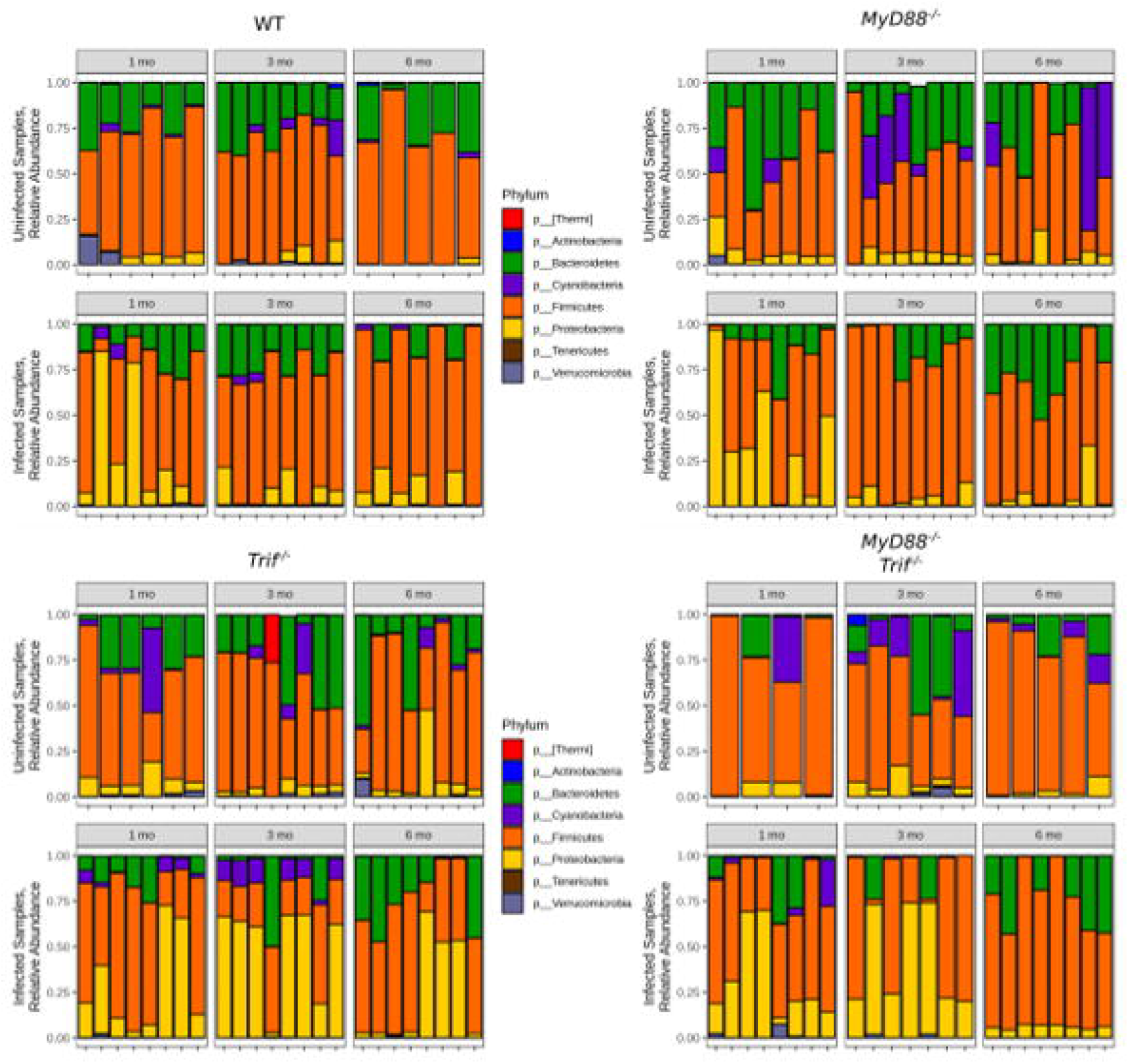
Relative abundance of different phyla across four genotypes. Samples are broken out by genotype, time, and infection status. Sequencing data processed in QIIME2, then plotted in PhyloSeq.

We observed predominance of 6 orders across all genotypes that include *Bacteroidales*, *Campylobacterales*, *Clostridiales*, *Lactobacillales*, *Streptophyta*, *Erysipelotrichales* (Figure 3). The order *Bacteroidales* was observed across all genotypes with no overall significant difference. *Clostridiales* was present in all genotypes but less abundant in DKO mice. However, it was observed that levels of *Clostridiales* dropped significantly in MyD88^−/−^ post infection as compared to uninfected samples especially at 3 months. *Campylobacterales* was observed more predominantly in infected samples in all four genotypes, as expected since *Helicobacter* is a member of the *Campylobacterales* order. However, the levels declined significantly at 3 and 6 months in MyD88^−/−^ mice and at 6 months in DKO mice although *Campylobacterales* can be seen throughout all time points in Trif^−/−^ infected mice. *Lactobacillales* was largely seen in MyD88^−/−^ infected mice especially at 3 months followed by DKO infected mice, WT infected mice and to a lesser extent Trif^−/−^ mice (Figure 3). *Streptophyta* was observed in uninfected samples of all genotypes at varying levels especially in MyD88^−/−^ mice but was completely lost post *H. felis* infection in MyD88^−/−^ mice at all time points and at 3 and 6 months in DKO mice. *Erysipelotrichales* was predominantly abundant at 6 months in Trif^−/−^ infected mice as compared to other genotypes. Apart from these orders *Rickettsiales* was observed mainly in uninfected MyD88^*−/−*^ and DKO mice as compared to Trif^−/−^ and WT mice and was almost lost upon infection with *H. felis*. In addition, another order *Bacillales* was observed in uninfected samples of Trif^−/−^, MyD88^−/−^ and WT but not in DKO uninfected mice. However, it was observed in DKO post infection in one sample both at 1 month and 3 months and in two samples in Trif^−/−^ at 6 months post infection with *H. felis*.

**Figure 3.**
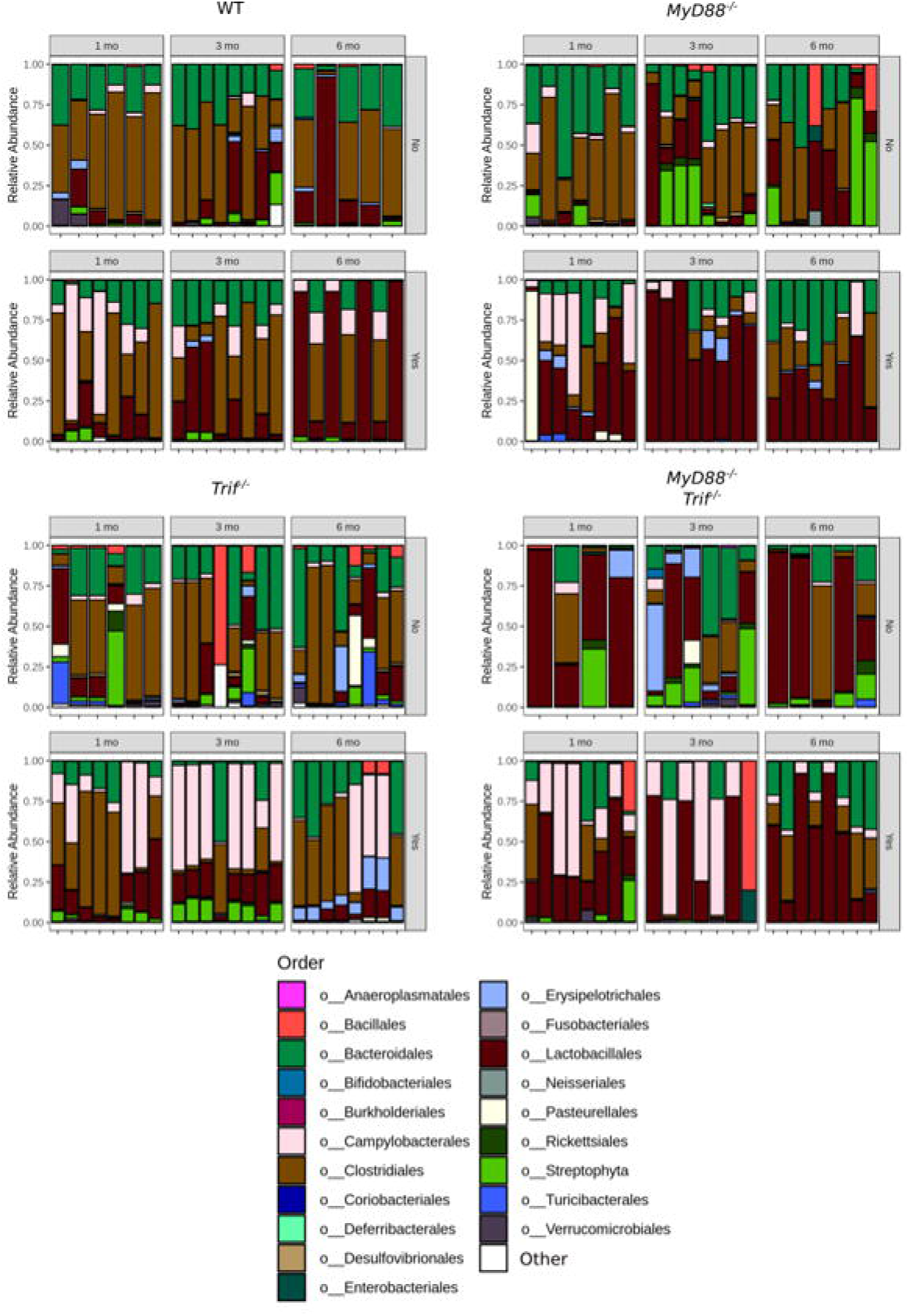
Relative abundance of different orders across four genotypes. Samples are broken out by genotype, time, and infection status. Sequencing data processed in QIIME2, then plotted in PhyloSeq.

### Association of *Lactobacillales* with infection and mouse genotype

Given the increase in *Lactobacillales* seen in infected mice with 16S sequencing (Figure 3) and the identification of *Lactobacillales* as a taxa driving community dissimilarity in the robust Aitchison distance (Figure 1), we further investigated the relationship between changes in *Lactobacillales* and disease progression. We used the Songbird (14) and Qurro (15) tools to analyze changes in *Lactobacillales* between conditions. Songbird calculates log-fold differentials of ratios of taxonomy units in each sample, a compositional analysis that accounts for absolute microbial abundance differences between samples. Qurro can then be used to visualize and compare these differentials between samples. This analysis revealed a highly significant increase in log fold differential between *Lactobacillales* and *Rickettsiales* (L/R) in infected communities compared to uninfected communities (Figure 4). Breaking this shift down by genotype showed the L/R differential was significantly increased in infected MyD88^−/−^ and DKO mice compared to uninfected. The ratio was not significantly different between infected and uninfected communities in Trif^−/−^ and WT mice. To confirm this finding, we repeated the analysis comparing *Lactobacillales* to other organisms present in the communities. Similar results were observed between *Lactobacillales* and *Streptophyta* and *Bacteroidales* again in MyD88^−/−^ and DKO as compared to Trif^−/−^ and WT mice (Figure 4). These findings indicate *Lactobacillales* is increased in infected communities in Myd88^−/−^ and DKO, but not WT and Trif^−/−^, genotypes.

**Figure 4.**
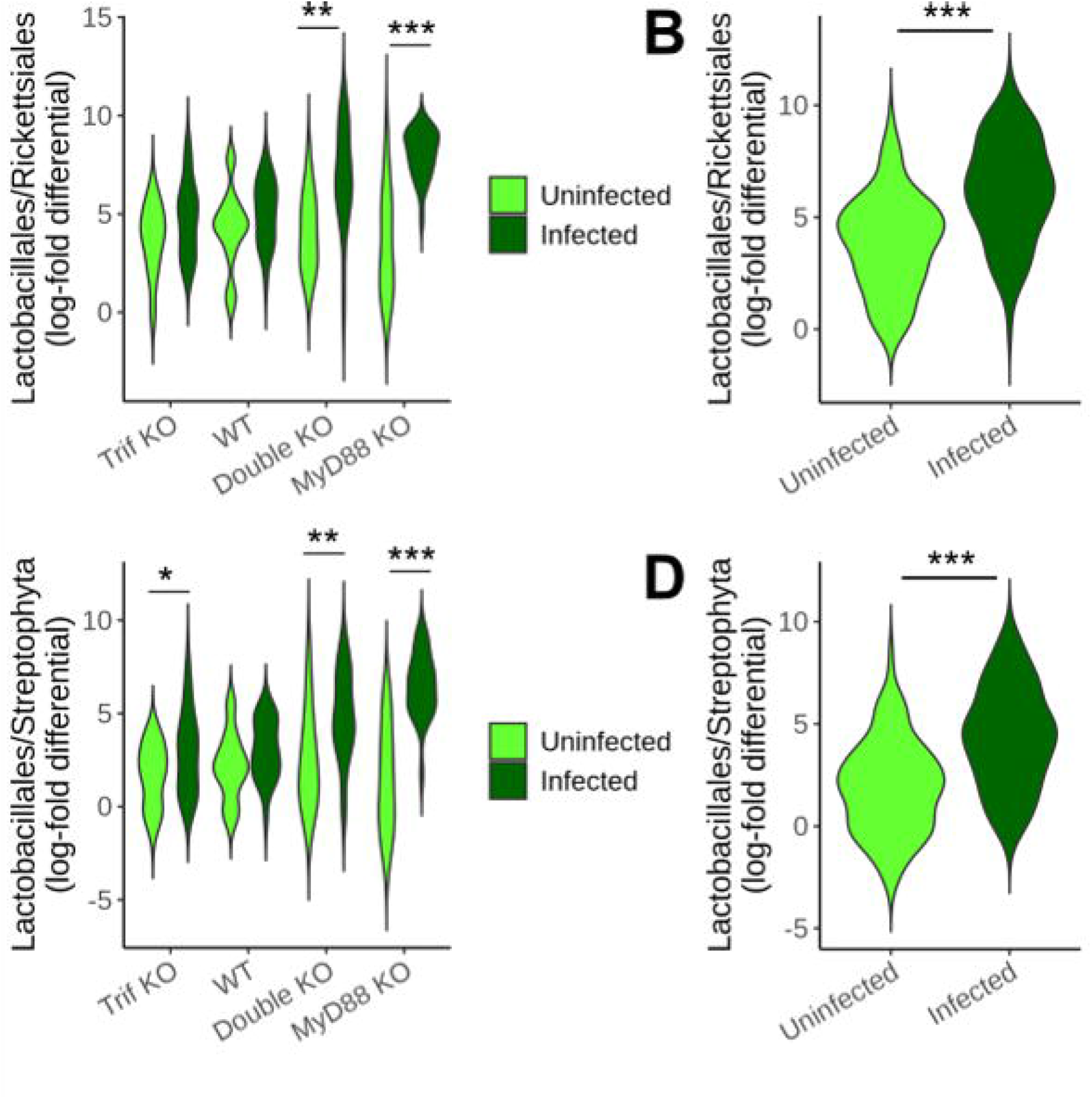
Log fold-differentials of ratio between different species across four genotypes. Log-fold differentials of the *Lactobacillales/Rickettsiales* (A, B) and *Lactobacillales/Streptophyta* (C, D) ratios between samples. A and C show ratios by genotype infection and status. B and D show ratios by infection status, all genotypes combined. Log fold-differentials were calculated and processed using Songbird and Qurro. Statistical significance determined by ANOVA (* p<0.05, ** p<0.01, *** p<0.005).

### *Lactobacillales* and disease progression

We next examined if there is an association between *Lactobacillales* ratios and gastric disease progression. Histological analysis revealed that Myd88^−/−^ genotype displays the worst gastric disease following infection, followed by the DKO genotype (Figure 5). Since these genotypes also possessed the highest levels of *Lactobacillales*, we hypothesized higher *Lactobacillales* ratios would be associated with higher mucous histology scores. Analysis of the L/R and L/S ratio and the mucous histology score of each sample with ordinal logistic regression demonstrated that mice with a higher gastric *Lactobacillales* ratio had a significantly higher likelihood of a higher histology metaplasia score. Given the association of *Lactobacialles* ratios and mucous metaplasia, we examined the ability of the *Lactobacillales* ratio to predict if a mouse had been infected with *H. felis*. We constructed receiver operator characteristic (ROC) curves for log-fold differentials of the orders *Campylobacterales, Lactobacillales*, and *Clostridiales* compared to *Rickettsiales* or *Streptophyta* (Figure 6). *Lactobacillales* log-fold differentials predicted mouse infection status at a rate similar to *Campylobacterales*, the order containing *Helicobacter* and the logical “gold standard” for infection predication. In comparison*, Clostridiales* log-fold differentials did not predict infection status at a rate better than random. Together, these data indicate *Lactobacillales* correlates strongly with Helicobacter infection status and mucous histology score. This association persists across genotypes, despite the differences observed in gastric microbiome composition between genotypes.

**Figure 5.**
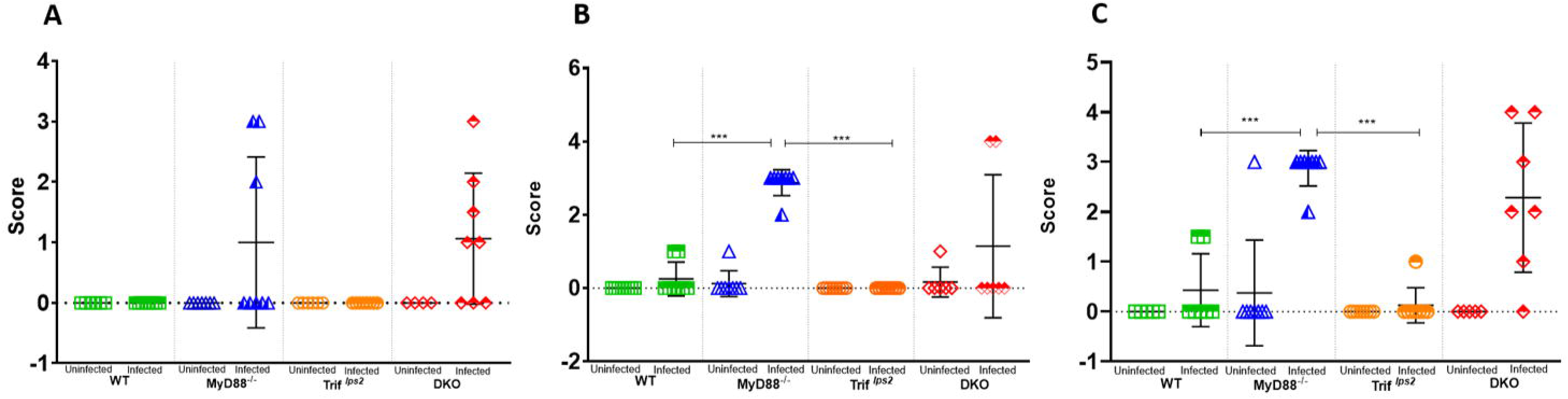
Histopathological scoring for Mucous Metaplasia. Following infection with *H.felis* for 1 month (A), 3 months (B) and 6 months (C), H&E-stained stomach sections from each mouse (WT and Myd88^−/−^, Trif^−/−^ and DKO) were evaluated for indications of pathology, and the mucous metaplasia was scored by a blinded comparative pathologist according to the criteria described in Materials and Methods. A P value of 0.05 was considered statistically significant. (A) 1 month post infection, n 14 for WT and n 15 for Myd88^−/−^, n 14 for Trif^−/−^ and n 12 for DKO mice; (B) 3 months post infection, n 16 for WT and n 16 for Myd88^−/−^, n 16 for Trif^−/−^ and n 13 for DKO mice; 6 months post infection, n 12 for WT and n 16 for Myd88^−/−^, n 16 for Trif^−/−^ and n 12 for DKO mice. (* p<0.05, ** p<0.01, *** p<0.005).

**Figure 6.**
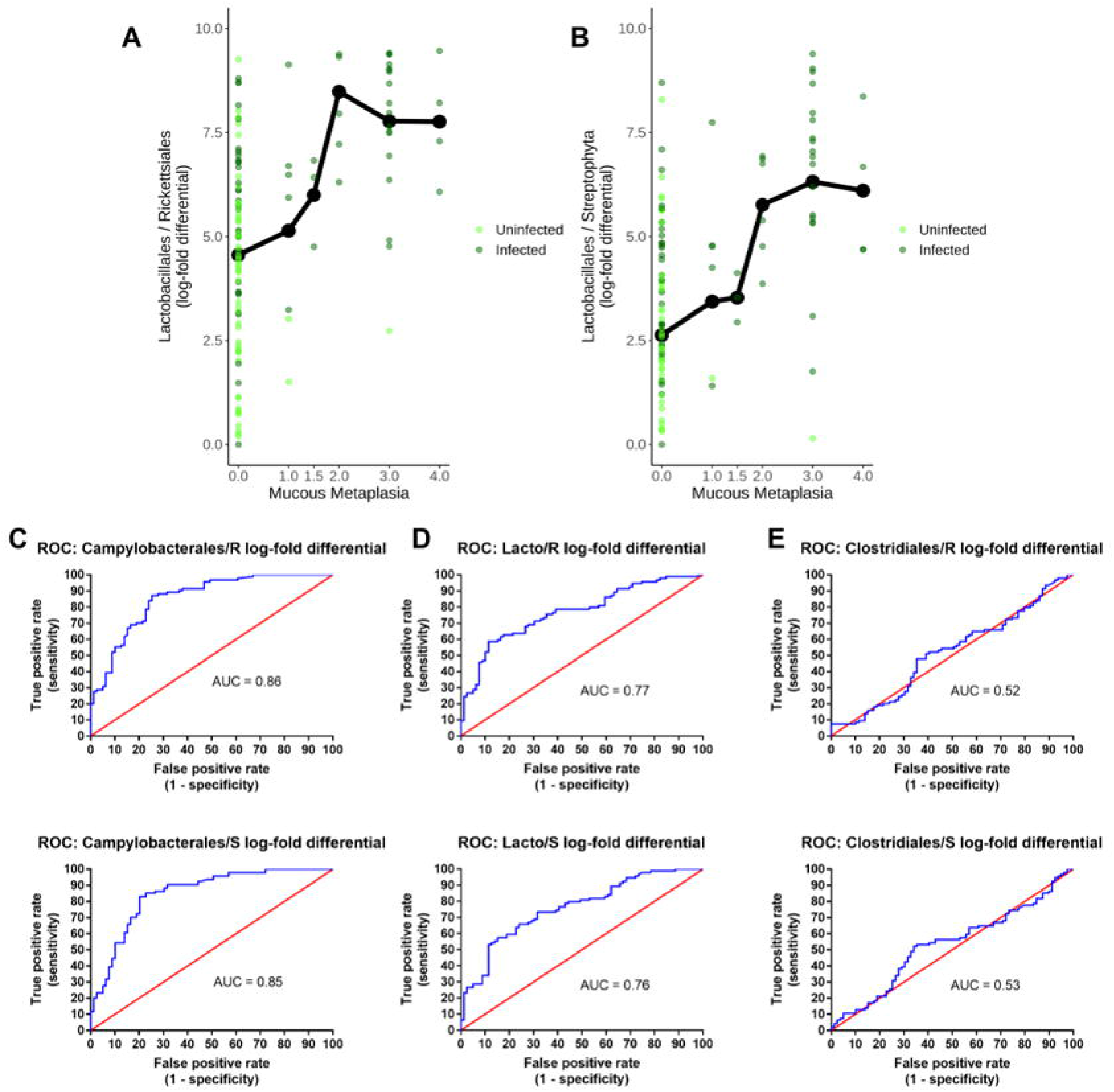
Predictive relationship between *Lactobacillales* and *Helicobacter* infection. A, B) Ordinal logistic regression analysis of log-fold differentials of *Lactobacillales/Rickettsiales* (A) and *Lactobacillales/Streptophyta* (B) ratios and gastric mucous histology score. The black line represents the average log-fold differential for each histology score. Ordinal logistic regression was calculated using R. C, D, E) ROC curve for log-differentials of *Campylobacterales*, *Lactobacillales*, and *Clostridiales*. R = *Rickettsiales*; S = *Streptophyta*. The blue line represents the performance of each ratio log fold-differential in predicting *Helicobacter* infection. The red line represents the result expected for a metric with a 50% chance of predicting infection. ROC plots were constructed in Prism7 using log fold-differentials from Songbird and Qurro.

## Discussion

Advancement in research on gut microbiome has discovered the existence of a diverse microbiome in the human gut in a delicate balance. The microbiota is vital for maintenance of human health and play an important role in energy metabolism, nutrient absorption and defense against pathogens (3, 18–20). However, if this balance is altered dysbiosis can lead to susceptibility to gastrointestinal pathogenesis and cancer.

The acidic environment of the stomach inhabits a smaller number of bacteria as compared to other parts of the gut, but dysbiosis due to various factors, genetic, environmental or pathogen invasion can lead to gastric cancer (3). The association of *Helicobacter* with gastric cancer has been well established, as it is characterized as a Type I carcinogen by the WHO (2). Previous studies have shown that *H. pylori* infections lead to an overall decrease in microbial density (21). Our findings indicate there is also an overall decrease in microbiome diversity upon *H. felis* infections across all genotypes.

In this study we sampled the mouse stomach mucosal tissue instead of fecal samples in order to understand the change in microbial diversity in different genotypes, with respect to time as well as infection status. Analyzing stomach mucosal tissue provides an accurate image of microbial activity during gastric cancer progression than fecal samples. Moreover, as previously described MyD88^−/−^ mice serve as fast progressing gastric cancer model, where gastric adenocarcinoma is reached within 6 months of infection with *Helicobacter* (6). Therefore, analyzing gastric mucosa for the periods of 1 month, 3 months and 6 months would provide a clear picture of microbial density fluctuations as gastric cancer progresses.

Previous studies in INS-GAS mice have shown that mice harboring a complex microbiome develop gastric cancer in 7 months post infection with *H. pylori* as compared to *H. pylori* infections in germ free mice where development of gastric cancer is prolonged. In addition, supplementation of germ-free mice with *Lactobacillus*, *Clostridium* species and *Bacteroides* species was sufficient to promote development of gastric cancer (21). This suggests the role of certain species in the gut microbiota in promoting gastric cancer progression. A study carried out on gastric cancer patients from high risk groups of Singapore and Malaysia population revealed a high relative abundance of lactic acid producing bacteria like *Lactococcus* and *Lactobacillus*, and also of oral cavity bacteria including *Fusobacterium*, *Veillonella*, *Leptotrichia*, *Haemophilus* and *Campylobacter* (22). To the contrary in our study we did not observe any increase in relative abundance of these bacteria except for *Lactobacilliales* and *Campylobacteriales*.

Comparing the gastric microbiome from Myd88^−/−^ mice to WT, Trif^−/−^ and DKO mice, we were able to intensively analyze how changes in the gastric microbiome could be connected to gastric cancer development and progression. In our study *Campylobacterales*, was observed in infected mice across all genotypes, which is expected as *Helicobacter* belongs to the order *Campylobacterales*. However, in MyD88^−/−^, our fast progressing gastric cancer model, we observed a reduction in *Campylobacterales* abundance at 3 months and 6 months. This could be attributed to the fact that with the advancement of gastric cancer lesions, increase in atrophy and decrease in acid secretion, there is a decrease in *Helicobacter* colonization, thus facilitating the increase in abundance of other bacteria. In accordance to our study, similar observation was seen in patients with advanced atrophic gastritis previous studies where low abundance of *H. pylori* led to hypo-chlorohydric stomachs, and the microbiota of gastric patients was dominated by other species (23, 24). Moreover, a study carried out on 273 gastric biopsies revealed no relationship between *H pylori* density and chronic gastritis (25). In contrast, studies carried out on human subjects by Basir et al., (26) revealed that increase in *H. pylori* colonization showed high correlation with severe chronic gastritis. Similar correlations were observed by Sayin S and Yakoob et al., (27, 28) in their respective studies on gastric patients. Thus, conflicting results have been observed when correlating helicobacter density to severity of disease.

The low abundance of *H. pylori* facilitates the dominance of other organisms of the microbiome. We observed an increase in *Lactobacilliales* in infected MyD88^−/−^ mice and DKO mice at 3 and 6 months. Previous studies on gastric cancer patients also showed increased abundance of *Lactobacilliales*, thus, suggesting its possible role in gastric cancer progression (23, 29) and in our case in fast progressing form of gastric cancer. Even though *Lactobacillus* species are utilized in probiotics and said to be beneficial for the host but high levels of lactic acid can be detrimental in case of gastric cancer. Lactate can serve has source of energy for tumor cells which can lead to increase ATP promoting inflammation (22, 30–33).

Furthermore, the *Lactobacillus* ratio comparison with histological data through ordinal logistic regression showed a significant correlation between *Lactobacillales* ratios and disease progression, while ROC curve analysis showed the ability of *Lactobacillales* to predict *Helicobacter* infection. These data all indicate *Lactobacillales* is closely associated with *Helicobacter* infection and gastric cancer progression.

On the other hand, we observed a complete loss of *Cyanobacteria*, at phylum level and *Streptophyta* at order level, in MyD88^−/−^ infected mice and DKO infected mice, but was present in Trif^−/−^ infected mice. *Cyanobacteria* and *Streptophyta* are usually not included in microbiome studies because they are food contaminants (34). But with their presence in Trif^−/−^ infected mice and absence in MyD88^−/−^ and DKO Infected mice, even when all the mice were housed in the same facility and fed the same food suggests that *Streptophyta* may have a role in delaying gastric cancer progression.

Recent studies in Taiwan, have shown that gastric cancer patients show increased colonization of *Clostridium* and *Fusobacterium* (35). Other studies have shown that INS-GAS germ free mice when supplemented with *Lactobacillus sp*., *Clostridium sp*., and *Bacteroides sp.*, develop gastric cancer (20). To the contrary, in our study the levels of *Clostriadales* significantly dropped in our fast progressing gastric cancer model, MyD88^−/−^ infected mice as compared to other genotypes, suggesting that even though *Clostriadales* may have a role in gastric cancer progression but not in a fast progressing gastric cancer model.

In conclusion, it can be suggested from our study that difference in genotypes could define the stomach microbiome diversity and make it more susceptible to development of gastric cancer post *Helicobacter* infections. Moreover, with the progression of disease the colonization of more *Lactobacillales* could contribute to faster development of gastric cancer. The presence of *Lactobacillales* and other taxa such as *Streptophyta* in the gastric microbiome needs to be further investigated in order to understand its probable role in fast progressing form of gastric cancer and stomach microbiome as a whole.

## Acknowledgements

This work is supported by the National Cancer Institute of the National Institute of Health under award R21CA210227. We thank S. Akira (Osaka University, Osaka, Japan) for the generous gift of Myd88^−/−^ original breeder mice.

## Abbreviations

*H. pylori*: *Helicobacter pylori*
WHO: World Health Organization
Myd88^−/−^: Myeloid differentiation primary response 88- deficient
*H. felis*: *Helicobacter felis*
Trif^−/−^: TIR-domain-containing adapter-inducing interferon-β
WT: Wild Type
DKO: double knockout
BHI: Brain Heart Infusion
QIIME2: Quantitative Insights into Microbial Ecology
OTU: Operational taxonomic unit

## Author Contributions

All listed authors have made an impactful and substantial contribution to this work. All authors have approved the final manuscript for publication.

