## Supplementary Figure Legend for "Microbiome signatures in a fast and slow progressing gastric cancer murine model and their contribution to gastric carcinogenesis"

### Supplementary Figure Legends

**Supplementary Figure 1**. **Relative abundance of different phyla across four genotypes.** Relative abundance for phyla plotted individually, broken out by genotype, time, and infection status. Plots were constructed in R with ggplot2, using sequencing counts from QIIME2 analysis.
