## Supplementary figures and images for "Microbiome signatures in a fast and slow progressing gastric cancer murine model and their contribution to gastric carcinogenesis"

### Supplementary Figure 1

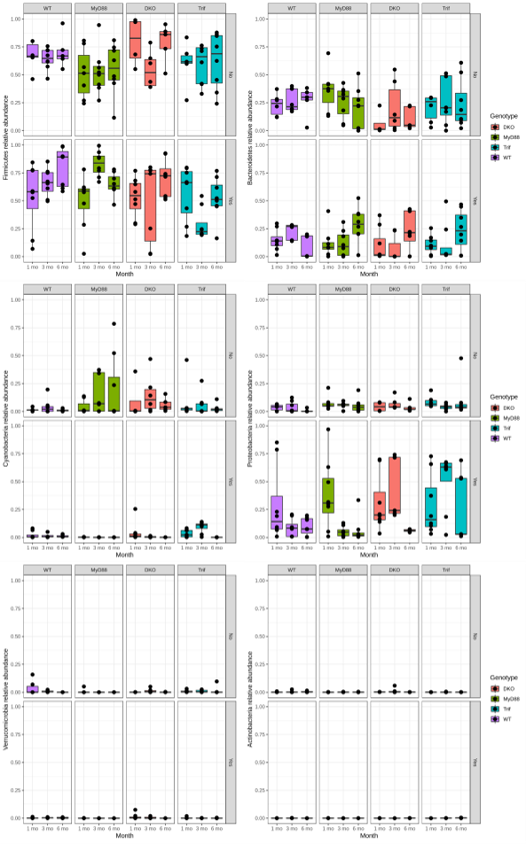
